# Complete diagnosis of leptospirosis in tropical reproductive cattle

**DOI:** 10.1101/327726

**Authors:** Gabriela Pacheco Sánchez, Fabio Almeida de Lemos, Mirian Dos Santos Paixão-Marques, Maria Fernanda Alves-Martin, Lívia Maísa Guiraldin, Wesley José Santos, Simone Baldini Lucheis

## Abstract

Leptospirosis is a worlwide zoonosis of great impact in both animal and public health. Bovine leptospirosis is commonly manifested by reproductive disorders, such as abortion, stillbirth and infertility; causing depletion of the economic balance of livestock farms, along with representing a health risk problem for farm workers. In view of these consequences, we aimed to evaluate the sanitary status of tropical cattle and their role as reproductive disseminators of leptospirosis. We analyzed blood and semen samples from 11 brazilian herds by three diagnostic methods -Culture, Microscopic Agglutination Test and Polymerase Chain Reaction. All animals were negative for bacteriological culture in Fletcher’s semisolid medium; 66% (264/400) animals were seropositive to at least one of 19 serovars (17 serogroups) of *Leptospira* spp. by MAT, given that 42.4% and 5.3% of animals presented titers against brazilians isolates Guaricura and Nupezo, respectively; furthermore, five animals were positive by PCR in blood and/or culture samples and three semen samples were positive by PCR (one of them also seropositive). These results highlight the coexistence of both disease's stages (acute and chronic) in the same environment, thus alert for venereal dissemination of leptospirosis, aggravating their sanitary condition and fomenting economic losses. We, authors, recommend the adoption of prophylactic measures, such as systemic vaccination, treatment of animals and improvement of hygienic-sanitary conditions.

## Introduction

Among reproductive diseases that affect livestock, leptospirosis has a high degree of importance, specially in tropical countries (1). This zoonosis is caused by members of genus *Leptospira* spp., which currently includes 22 species and more than 300 serovars (2). Bovine leptospirosis represents an animal health problem, given its common manifestation like chronic reproductive disorders, such as abortion, stillbirth and infertility (3,4). Clinical signs of the acute presentation and mortality are most frequent in calves (5). Furthermore, leptospirosis also represents a public health risk for farm workers along with decrease in economic balance of livestock farms (6). For diagnosis of leptospirosis, many current options are avaliable depending on sample type and clinical phase of disease, each method has its own advantages and disadvantages. Microscopic aglutination test - MAT, remains the reference test for routine diagnosis (7,8). Concordance among positive MAT results and positive outcomes by other techniques has proved that MAT is an efficient diagnostic test for prediction of infecting serogroup (9); althought, another study proved that cattle may not react against their own isolate by MAT, demanding caution when evaluating negative MAT results for carrier status (10). Countries with strong livestock production, like Brazil, have a particular concern on elucidate important features regarding leptospirosis. Many research groups aim to explain the dissemination and risk factors of this disease (11). Throughout the years, the interest on understanding leptospirosis pathogenesis has led to studies envolving possible transmission routes. Venereal transmission has attracted attention since the report of genital Hardjo infection in naturally infected cattle (12), and presence of *Leptospira* spp. was demonstrated in bovine semen by PCR (13), proving that the bacteria can also be established in gonads, suggesting possible sexual transmission. These informations should be considere when assessing disease’s control programs, along with endemic serovars, antibiotic and vaccination availability, besides epidemiological studies (14,15). In this context, we aimed to evaluate the sanitary status of this zoonosis in clinically asymptomatic cattle from a tropical region of Brazil.

## Material and Methods

The area chosen for this study was the microregion of Bauru, which is a country town of São Paulo, Brazil. Bauru is a tropical area known for its hot temperature, typically from 59 F to 86 F; with extreme seasonal variation in rainfall, the least rain falls in winter and the most rain falls around summer, encompassing up to 12 inches.

In order to estimate sample size for this study, we used the online program Epi Info^TD^ (http://www.openepi.com), with confidence interval of 95% and based on a preview study that found prevalence of 58.7% in the city of Bauru, years before (16). Sample size calculated for disease frequency was 373 samples.

We analyzed blood samples of 11 herds from five farms (representing 400 animals); as to semen samples from two herds. Cattle were raised for reproductive purposes and shared water fountain and pastures among most herds studied. All farms presented historic of reproductive failures and no vaccination program installed for leptospirosis. Animals studied were clinically asymptomatic for leptospirosis. Approximately 5-10 ml of blood samples were collected using sterile syringes, by caudal tail vein venipuncture, and evacuated into one tube with anticoagulant and one without it. Semen samples were collected by electroejaculation (Electro-Ejaculator DUBOI), approximately 2 - 5 ml of semen were obtained from each animal. All samples were identified, refrigerated and transported to the Animal Sanity Laboratory of Paulista Agency of Agribusiness Technology, APTA, Bauru, Brazil. At the laboratory, blood and semen samples were inoculated in Fletcher medium for bacteriological isolation, employing a serial dilution technique with some modifications (17). From all dilutions, 0.5 mL were inoculated into a tube containing semisolid Fletcher’s culture medium (Difco^®^) with 0.15% agar, supplemented with 100 μg of 5-fluorouracil/mL and 1% sterile rabbit serum and inactivated at 56 °C for 30 min. Cultures were incubated at 28-30 °C for 16 weeks. Dark-field microscopy evaluation of tubes was performed every two weeks. For detection of anti-*Leptospira* antibodies, Microscopic Agglutination Test (MAT) was used with a panel of 19 serovars (including two native isolates) representing 17 pathogenic and intermediate serogroups (table 1), according to international standards (18). Serum samples were considered reactive when reached titers ≥ 100 and the ultimate reactive serogroup was determined by election of the highest titer presented; when presence of coagglutinations, all serovars involved were considered as positive.

**Table 1.**
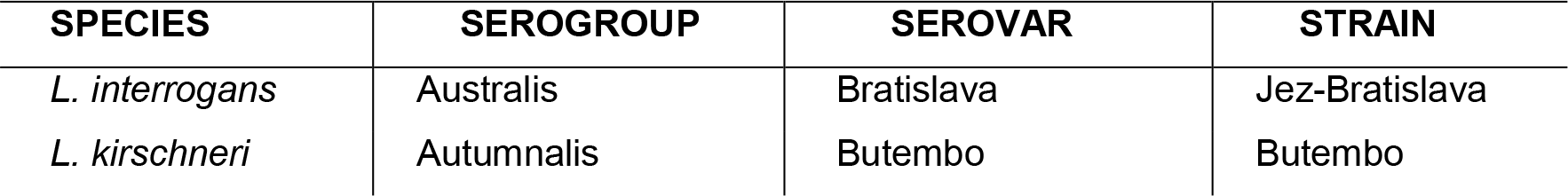
Serovars used in Microscopic Agglutination Test (MAT).

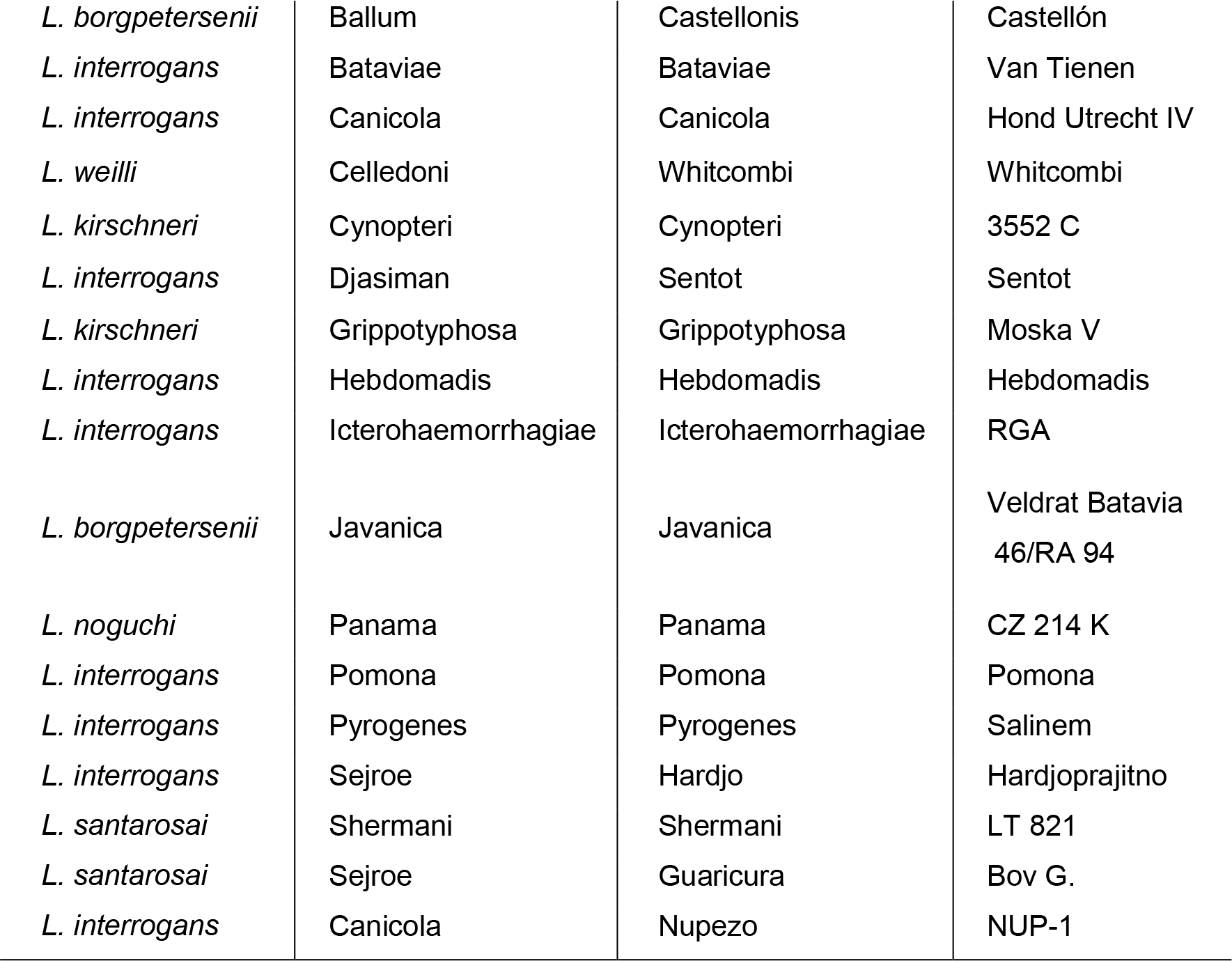

Serum samples were considered reactive when reached titers ≥ 100.

DNA from blood and culture samples (for confirmatory purposes) was extracted for Polymerase Chain Reaction (PCR) using the Illustra Blood Genomic Prep Mini Spin kit (GE Healthcare^®^), while DNA from semen samples was extracted using DNAzol^®^ reagent, both according to manufacturers’ recommendations. Primers employed amplify a fragment of 331 base pairs length (19): LEP 1 (5′ GGCGGCGCGTCTTAAACATG 3′) and LEP 2 (5′ TTCCCCCCATTGAGCAAGATT 3′). The final reaction volume was 25 μL, including 2.5 μL of PCR buffer solution (50 mM KCl and 10 mM Tris-HCl, pH 8.0), 0.75 μL of MgCl_2_ (1.5 mM), 0.5 μL of dNTP solution (0.2 mM), 0.5 μL of ***Taq Platinum*** DNA (1 U) (Invitrogen^®^), 0.5 μL of each primer (10 pM), 17.75 μL of ultrapure water, and 2 μL of the DNA extracted from each sample. PCR reaction was conducted in a Mastercycler^®^ gradient thermal cycler (Eppendorf), according to the protocol described by Merien et al., 1992, with modifications. The amplified products were visualized by electrophoresis in 1.5% agarose. Cultures of *Leptospira interrogans* serovar Copenhageni in EMJH medium were used as positive controls, as well as ultrapure water like negative controls.

## Results

Cultures of all samples were performed to detect presence of *Leptospira* spp. in blood and semen samples. All samples were negative to bacteriological culture, using visualization by dark-field microscope. All eleven herds presented titers against at least one pathogen serogroup of *Leptospira* spp. (minimum of 50% animals reactive per herd); 66% (264/400) of animals were positive to MAT. Most reactive serogroups were Sejroe, serovar Guaricura (local strain) with 112 positive animals and serovar Hardjo with 102 positive animals; and serogroup Autumnalis with 77 positive MAT reactive samples. Highest titers reached 3200 when tested against serovar Guaricura and serovar Hardjo. Coaglutinations reached up to 37.1% of all MAT positive outcomes. Analysis by PCR of blood samples showed some controversial results. Out of all negative blood cultures, four animals turned out to be positive by confirmatory culture PCR. When analyzing direct blood PCR results, we noted that three animals were positive for presence of *Leptospira* spp.; perturbation aroused when confirming that out of the positive samples, only one was positive both in confirmatory culture PCR as to in blood PCR. Thus, confirming the conflicted scenario of having the same theoretical sample positive in one test and negative in the other. None of these positive animals had MAT positive outcomes (table 2).

**Table 2.**
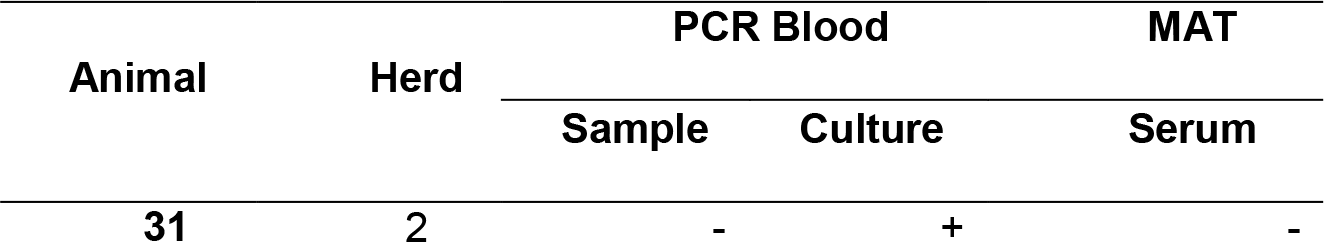
Molecular and serological outcome for bulls positive by PCR in semen and/or blood samples.

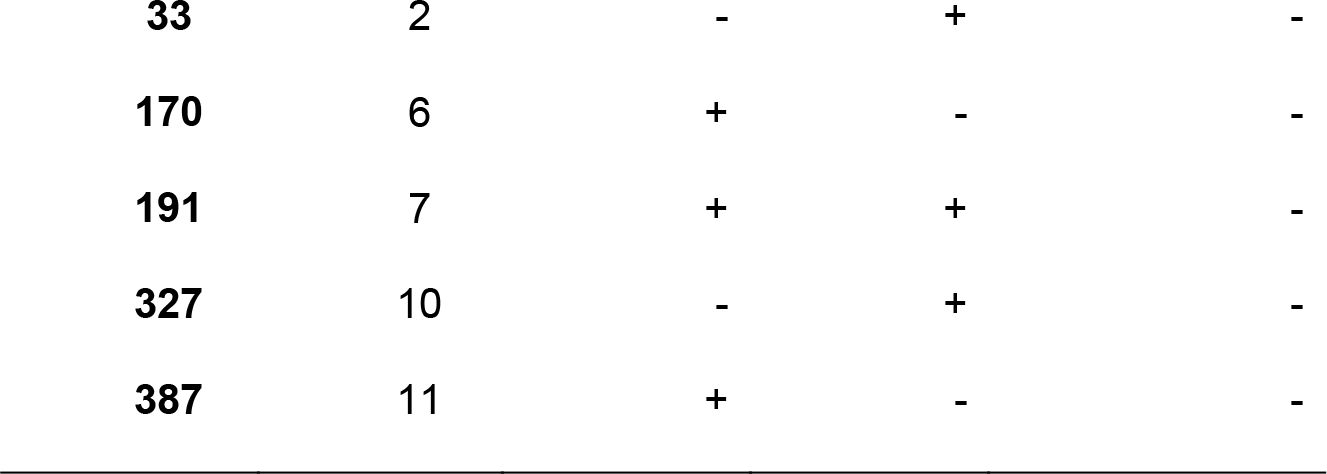

Analysis by PCR of semen displayed some interesting results as well. Two bulls were positive for confirmatory PCR of negative culture (dark-field microscopy); three bulls were positive for semen PCR and for semen culture confirmation PCR; and other two bulls were positive solely for culture confirmation PCR, but not for direct semen PCR. Among these positive bulls, three were also positive by MAT (table 3).

**Table 3.** Molecular and serological outcome for bulls positive by PCR in semen and/or blood samples.

| Animal | Herd | PCR Blood |  | PCR Semen |  | MAT |
| --- | --- | --- | --- | --- | --- | --- |
|  |  | Sample | Culture | Sample | Culture | Serum |
| **5** | 1 | - | - | - | + | + |
| **7** | 1 | - | - | + | + | - |
| **27** | 2 | - | - | + | + | - |
| **29** | 2 | - | - | - | + | + |
| **30** | 2 | - | - | + | + | + |
| **31** | 2 | - | + | - | - | - |
| **33** | 2 | - | + | - | - | - |

## Discussion

In order to evaluate sanitary state of the disease and not just merely exposure to the bacteria, we used serovars from pathogenic species of *Leptospira* spp., one serovar representative from each serogroup and also two native brazilian isolates, serovar Guaricura (*Leptospira santarosai* serogroup Sejroe) and serovar Nupezo (*Leptospira canicola* serogroup Canicola). Serovar Nupezo showed reactivity when compared to serovar Canicola, supporting the inclusion of local strains for more sensitive MAT results. Serogroup Sejroe had most positive outcomes, however serogroups Autumnalis and Hebdomadis were also quite reactive, suggesting possible contact among cattle with wild animals, given that only serogroup Sejroe is known to be adaptive to cattle and the other two serogroups have been reported in wild animals (20,21). Several studies around Brazilian territory showed the magnitude and importance of leptospirosis in this tropical region, and despite variety of these studies’ outcomes, the seriousness can’t be disguised (table 4).

**Table 4.** Prevalence studies about bovine leptospirosis in Brazil.

| Region | State | Prevalence | Animals | Frequent serogroups | References |
| --- | --- | --- | --- | --- | --- |
| North | Maranhão | 35.94% | 4832 cows | Sejroe | (22) |
| Northeast | Bahía | 45.42% | 10 823 bovines | Sejroe | (23) |
| Central-west | Mato Grosso do Sul | 98.8% | 1801 cows | Sejroe | (24) |
| Southeast | Santa Catarina | 65.53% | 3945 cows | Autumnalis and Sejroe | (25) |
| South | Rio de Janeiro | 14% | 120 cows | Sejroe and Shermani | (26) |
|  | São Paulo | 45.8% | 2761 bovines | Sejroe | (16) |

Considering that 66% (264/400) of cattle were seropositive, furthermore six animals were positive for PCR of blood and/or culture of blood, as three bulls were also positive by confirmatory PCR of semen cultures (one of them confirmed by semen PCR as well), we can suggest there is an imminent present exposure of cattle to the disease in those farms, the same ones that constitute a representation of the extensive livestock production system in this tropical region.

PCR outcomes also showed trustworthiness when compared to culture, given that results that were negative by culture were positive by PCR, which was no surprise considering difficultness of visualization of spirochetes by dark-field microscopy (7). Surprise came when comparing PCR results between negative cultures and samples, animals were positive on blood culture but not on whole blood or vice versa, suggesting that there could be an inhibitor in blood that was diluted when cultured. Similar results occurred with two semen samples, thus same explanation can be granted (27).

Some current studies have attempt to confirm presence of leptospiral DNA in reproductive secretions. In cattle, two research group evaluated semen of seropositive bulls with unsuccessful results (28,29)), but more recently another research group was capable to detect Leptospiral DNA from vaginal fluid of cows (30). Other studies have been executed in goats and sheeps, with positive results for vaginal fluids and semen (31); and in mares and horses, which lead to detection of leptospiral DNA in vaginal fluids, genital tract and semen (32,33). Thus, the results obtained in this study enhance current knowledge of not only sanitary status of the disease, but to possible participation of sexual transmission in farms representative of the extensive livestock production farming in this tropical region.

These farms do not have an established surveillance program for leptospirosis, therefore lack of prevention could be one of the main reasons for the serological and molecular results. We, authors, strongly recommend the adoption of prophylactic measures, such as systemic vaccination, treatment of animals and improvement of hygienic-sanitary conditions, given the results obtained.

## Acknowledgments

We will like to thank to National Council for Scientific and Technological Development - CNPq and Paulista Agency of Agribusiness Technology - APTA Bauru.

## References

1. Martins G, Lilenbaum W. The panorama of animal leptospirosis in Rio de Janeiro, Brazil, regarding the seroepidemiology of the infection in tropical regions. BMC Vet Res [Internet]. 2013;9:237. Available from: http://www.pubmedcentral.nih.gov/articlerender.fcgi?artid=4220826&tool=pmcentrez&rendertype=abstract

2. Picardeau M. Virulence of the zoonotic agent of leptospirosis: still terra incognita? Nat Rev Microbiol [Internet]. 2017 [cited 2017 Aug 23];15(5):297–307. Available from: http://doi.org/10.1038/nrmicro.2017.5

3. Ellis W. Animal Leptospirosis. In: Leptospira and Leptospirosis. Springer; 2015. p. 295.

4. Martins G, Penna B, Hamond C, Leite RC-K, Silva A, Ferreira A, et al. Leptospirosis as the most frequent infectious disease impairing productivity in small ruminants in Rio de Janeiro, Brazil. Trop Anim Health Prod [Internet]. 2012;44(4):773–7. Available from: http://www.ncbi.nlm.nih.gov/pubmed/21898182

5. Lilenbaum W, Martins G. Leptospirosis in cattle: A challenging scenario for the understanding of the epidemiology. Transbound Emerg Dis. 2014;61(SUPPL1.):63–8.

6. Langoni H, de Souza LC, da Silva a V, Cunha ELP, da Silva RC. Epidemiological aspects in leptospirosis. Research of anti-Leptospira spp antibodies, isolation and biomolecular research in bovines, rodents and workers in rural properties from Botucatu, SP, Brazil [Aspectos epidemiológicos nas leptospiroses: Pesquisa d. Brazilian J Vet Res Anim Sci [Internet]. 2008;45(3):190–9. Available from: http://www.scopus.com/inward/record.url?eid=2-s2.0-78149436726&partnerID=40&md5=e85cfb1ce0f76c5de43234e55c5a5b3b

7. Picardeau M. Diagnosis and Epidemiology of leptospirosis. Diagnostic épidémiologie la leptospirose [Internet]. 2013 [cited 2017 Aug 24];43:1–9. Available from: http://dx.doi.org/10.1016/j.medmal.2012.11.005

8. Picardeau M, Bertherat E, Jancloes M, Skouloudis AN, Durski K, Hartskeerl R a. Rapid tests for diagnosis of leptospirosis: current tools and emerging technologies. Diagn Microbiol Infect Dis [Internet]. 2014 Jan [cited 2014 Oct 22];78(1):1–8. Available from: http://www.ncbi.nlm.nih.gov/pubmed/24207075

9. Blanco RM, dos Santos LF, Galloway RL, Romero EC. Is the microagglutination test (MAT) good for predicting the infecting serogroup for leptospirosis in Brazil? Comp Immunol Microbiol Infect Dis. 2016;44:34–6.

10. Libonati H, Pinto PS, Lilenbaum W. Seronegativity of bovines face to their own recovered leptospiral isolates. Microb Pathog. 2017;108:101–3.

11. Fávero JF, Araújo HL De, Lilenbaum W, Machado G, Tonin AA, Baldissera MD, et al. Bovine leptospirosis: Prevalence, associated risk factors for infection and their cause-effect relation. Microb Pathog [Internet]. 2017 [cited 2017 Aug 22];107:149–54. Available from: http://dx.doi.org/10.1016/j.micpath.2017.03.032

12. Ellis W, Songer J, Montgomery J, Cassells J. Prevalence of Leptospira interrogans serovar hardjo in the genital and urinary tracts of non-pregnant cattle. Vet Rec. 1986;118(1):11–3.

13. Heinemann MB, Garcia JF, Nunes CM, Morais ZM, Gregori F, Cortez A, et al. Detection of leptospires in bovine semen by polymerase chain reaction. Aust Vet J. 1999;77(1):32–4.

14. Adler B, de la Peña Moctezuma A. Leptospira and leptospirosis. Vet Microbiol [Internet]. 2010 Jan 27 [cited 2014 May 24];140(3-4):287–96. Available from: http://www.ncbi.nlm.nih.gov/pubmed/19345023

15. Martins G, Lilenbaum W. Control of bovine leptospirosis: Aspects for consideration in a tropical environment. Res Vet Sci [Internet]. 2017 [cited 2017 Aug 24];112:156–60. Available from: http://dx.doi.org/10.1016/j.rvsc.2017.03.021

16. Langoni H, Meireles LR, Gottschalk S, Cabral KG, Silva AV. Perfil Sorológico Da Leptospirose Bovina Em Regiões Do Estado De São Paulo. Arq Inst Biol, São … [Internet]. 2000 [cited 2018 Apr 1];37–41. Available from: http://revistas.bvs-vet.org.br/arqib/article/download/25774/26642

17. Passos EDC, Vasconcellos SA, Ito FH, Yasuda PH, Junior RN. Isolamento de Leptospiras a partir do tecido renal de hamsters experimentalmente infectados com Leptospira interrogans sorotipo pomona. Emprego das técnicas da pipeta Pasteur e a das diluições seriadas em meio de cultura de Fletcher tratado com 5-Fluor-ur. Rev da Fac Med Vet e Zootec da Univ São Paulo. 1988;25(2):221–35.

18. OIE. Leptospirosis. In: Manual of Diagnostic Tests and Vaccines for Terrestrial Animals - Web Format. Fifth edit. Paris: OIE; 2014. p. 1–15.

19. Merien F, Amouriaux P, Perolat P, Baranton G, Girons I Saint. Polymerase Chain Reaction for Detection of Leptospira in Clinical Samples. J Clin Microbiol. 1992;30(9):2219–24.

20. Paixão MDS, Alves-Martin MF, Tenório MDS, Starke-Buzetti W a, Alves ML, da Silva DT, et al. Serology, isolation, and molecular detection of Leptospira spp. from the tissues and blood of rats captured in a wild animal preservation centre in Brazil. Prev Vet Med [Internet]. 2014 Jul 1 [cited 2014 Oct 23];115(1-2):69–73. Available from: http://www.ncbi.nlm.nih.gov/pubmed/24703251

21. Pinto PS, Pestana C, Medeiros MA, Lilenbaum W. concept of adaptability of leptospires to cattle. 2017;172(April):156–9.

22. Silva FJ, Conceição WLF, Fagliari JJ, Girio RJS, Dias R a., Borba MR, et al. Prevalência e fatores de risco de leptospirose bovina no estado do maranhão. Pesqui Vet Bras. 2012;32(4):303–12.

23. Oliveira FCS, Azevedo SS, Pinheiro SR, Viegas SARA, Batista CSA, Coelho CP, et al. Soroprevalência Da Leptospirose Em Fêmeas Bovinas Em Idade Reprodutiva No Estado De São Paulo, Brasil. Arq Inst Biol. 2009;76(4):539–46.

24. Figuereido A de O, Pellegrin AO, Gonçalves VSP, Freitas EB, Monteiro LARC, Oliveira JM de, et al. Prevalência e fatores de risco de leptospirose em bovinos de Mato Grosso do Sul. Pesqui Veterinária Bras. 2009;29(5):375–81.

25. Tonin AA, Azevedo MI De, Escobar TP, Casassola I, Gaspareto L, Schafer A, et al. LEPTOSPIROSE BOVINA: AUMENTO NA INCIDÊNCIA DA Leptospira interogans SOROVAR butembo NO REBANHO DO ESTADO DE SANTA CATARINA, BRASIL. Microbiol. 2010;4(4):294–7.

26. Folly M, Estadual U, Fluminense N. Anti-Leptospira agglutinins in farm workers, bovines and canines performed in the microregion of Itaperuna. … J Bras Ciência Anim. 2013;6(July 2013):406–17.

27. Alaeddini R. Forensic implications of PCR inhibition - A review. Forensic Sci Int Genet [Internet]. 2012 [cited 2017 Aug 24];6(3):297–305. Available from: http://dx.doi.org/10.1016/j.fsigen.2011.08.006%0A

28. Magajevski FS, Silva Girio RJ, Mathias LA, Myashiro S, Genovez MÉ, Scarcelli EP. Detection of Leptospira spp. in the semen and urine of bulls serologically reactive to Leptospira interrogans serovar Hardjo. Brazilian J Microbiol. 2005;36:43–7.

29. Vinodh R, Raj GD, Govindarajan R, Thiagarajan V. Detection of Leptospira and Brucella genomes in bovine semen using polymerase chain reaction. Trop Anim Health Prod [Internet]. 2008 [cited 2017 Aug 24];40:323–9. Available from: http://doi.org/10.1007/s11250-007-9110-5%0D

30. Loureiro AP, Pestana C, Medeiros MA, Lilenbaum W. High frequency of leptospiral vaginal carriers among slaughtered cows. Anim Reprod Sci [Internet]. 2017 [cited 2017 Aug 23];178:50–4. Available from: https://doi.org/10.1016/j.anireprosci.2017.01.008

31. Lilenbaum W, Varges R, Brandão FZ, Cortez A, de Souza SO, Brandão PE, et al. Detection of Leptospira spp. in semen and vaginal fluids of goats and sheep by polymerase chain reaction. Theriogenology [Internet]. 2008 [cited 2017 Aug 24];69(7):837–42. Available from: http://doi.org/10.1016/j.theriogenology.2007.10.027

32. Hamond C, Martins G, Medeiros MA, Lilenbaum W. Presence of leptospiral DNA in semen suggests venereal transmission in horses. J Equine Vet Sci [Internet]. 2013 [cited 2017 Aug 24];33:1157–9. Available from: http://dx.doi.org/10.1016/j.jevs.2013.03.185

33. Hamond C, Pestana CP, Rocha-de-Souza CM, Cunha LER, Brandão FZ, Medeiros MA, et al. Presence of leptospires on genital tract of mares with reproductive problems. Vet Microbiol [Internet]. 2015 [cited 2017 Aug 24];179:264–9. Available from: https://doi.org/10.1016Zj.vetmic.2015.06.014

